# Mapping Transcription Factor-Nucleosome Dynamics from Plasma cfDNA

**DOI:** 10.1101/2021.04.14.439883

**Authors:** Satyanarayan Rao, Amy L. Han, Alexis Zukowski, Etana Kopin, Carol A. Sartorius, Peter Kabos, Srinivas Ramachandran

## Abstract

Cell-free DNA (cfDNA) contains a composite map of the epigenomes of its cells-of-origin. Tissue-specific transcription factor (TF) binding inferred from cfDNA could enable us to track disease states in humans in a minimally invasive manner. Here, by enriching for short cfDNA fragments, we directly map TF footprints at single binding sites from plasma. We show that the enrichment of TF footprints in plasma reflects the binding strength of the TF in cfDNA tissue-of-origin. Based on this principle, we were able to identify the subset of genome-wide binding sites for selected TFs that leave TF-specific footprints in plasma. These footprints enabled us to not only identify the tissue-of-origin of cfDNA but also map the chromatin structure around the factor-bound sites in their cells-of-origin. To ask if we can use these plasma TF footprints to map cancer states, we first defined pure tumor TF signatures in plasma *in vivo* using estrogen receptor-positive (ER+) breast cancer xenografts. We found that the tumor-specific cfDNA protections of ER-α could distinguish WT, ER-amplified, and ER-mutated xenografts. Further, tumor-specific cfDNA protections of ER-α and FOXA1 reflect TF-specific accessibility across human breast tumors, demonstrating our ability to capture tumor TF binding in plasma. We then scored TF binding in human plasma samples and identified specific binding sites whose plasma TF protections can identify the presence of cancer and specifically breast cancer. Thus, plasma TF footprints enable minimally invasive mapping of the regulatory landscape of cancer in humans.

## Introduction

Transcription factors (TFs) are at the apex of gene regulation (*1, 2*). They usually bind small stretches of DNA in a sequence-specific manner (*3, 4*). The size of the mammalian genomes is several orders of magnitude greater than the size of TF binding motifs. Hence, there are many more transcription factor binding site (TFBS) sequences that occur by chance compared to functional TFBS (*5*). Although the question of how TFs discriminate functional binding sites from random motif occurrences is still actively investigated (*6-10*), at least two mechanisms enable us to connect TF binding to cell state. First, the cell type-specific expression of TFs restricts the pool of motifs recognized in a given cell type. Second, most motifs in the genome are occluded by nucleosomes most of the time (*11-15*). As a result, the sites in the genome bound by any given TF contribute to the epigenomic signature of a cell type. Furthermore, since functional TF binding drives gene regulation, mapping a TF’s binding sites in a cell also contributes to an understanding of the regulatory landscape of the cell (*16, 17*). Methods like Chromatin immunoprecipitation with DNA sequencing (ChIP-seq), chromatin immunoprecipitation, exonuclease digestion and DNA sequencing (ChIP-exo) and Cleavage Under Target & Release Using Nuclease (CUT&RUN) have been used to identify binding sites of human TFs across cell-types (*18-21*). Here, we show how to leverage this vast knowledge of TF binding in different cell types to map TF footprints in human plasma.

Dying cells in the human body release their content into the bloodstream (*22*). Genomic DNA that is bound by nucleosomes and TFs escapes endogenous nucleases and so remains protected in plasma (**Figure 1A**, (*23*)). Fragmentomics seeks to uncover tissue-of-origin of cfDNA using the information in cfDNA fragment length. Fragmentomics had its earliest application in prenatal diagnosis and is now being explored as an alternative to mutations and methylation profiling to identify cfDNA tissue-of-origin in cancer (*24-26*). cfDNA properties such as promoter nucleosome dynamics, locus-specific fragment length distribution, nucleosome-spacing in gene bodies, and nucleosome depletion at promoters have been used to identify tissue-of-origin of cfDNA in order to aid detection of cancer (*23, 27, 28*). Since TFs and nucleosomes protect distinctly different lengths of DNA, cfDNA facilitates direct mapping of protein-DNA interactions in their cells-of-origin (*23*). TF binding from cfDNA has also been characterized by averaging across thousands of putative sites, either looking at short protections (*23*) or by inferring TF binding by nucleosome depletion at TFBS (*29*).

**Figure 1.**
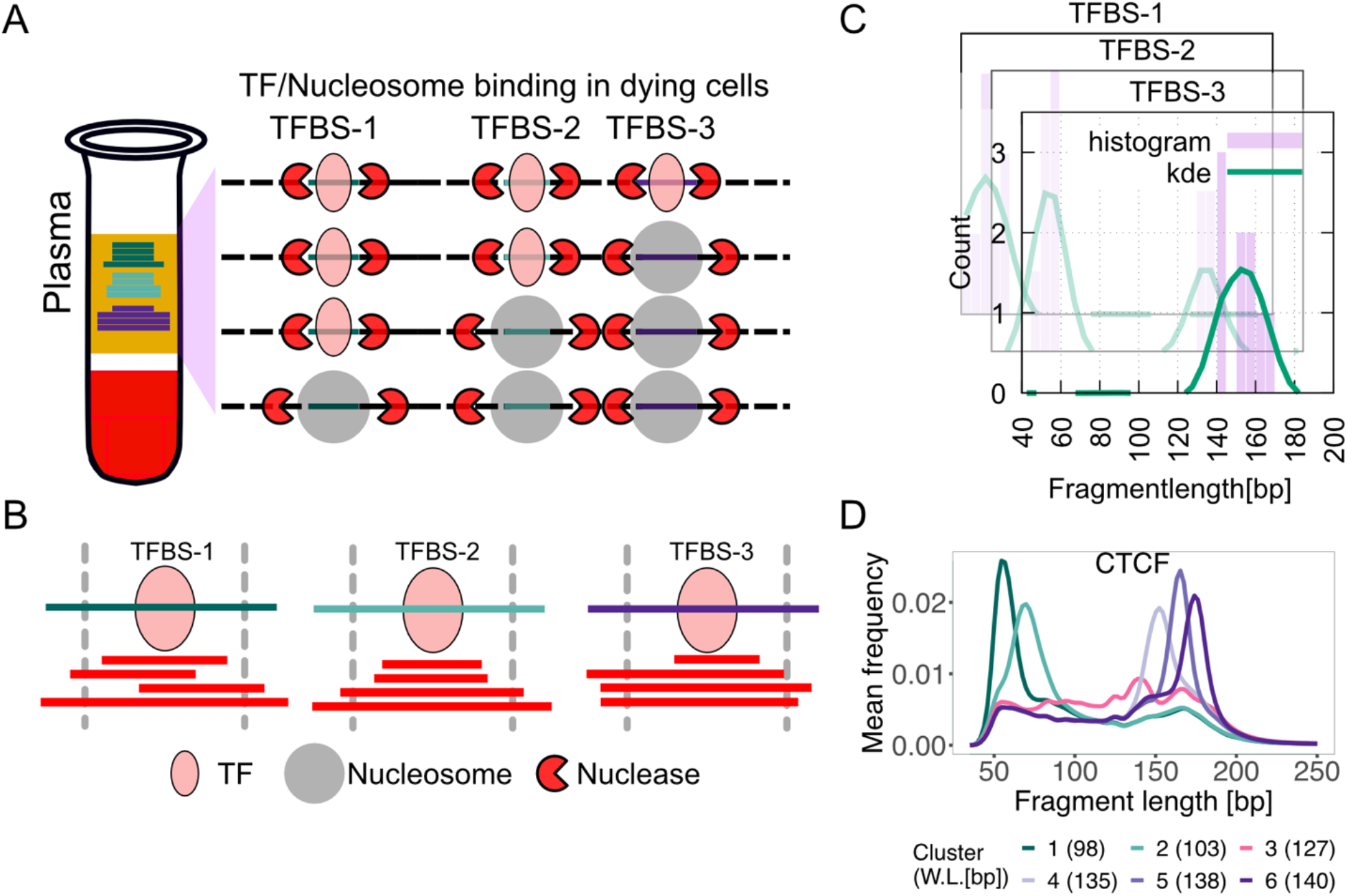
Schematic for identifying subset of binding sites with TF footprints. **A**) When TFs or nucleosomes are bound at TF binding sites, they protect different lengths of DNA from nucleases in dying cells in the human body. **B**) When sequenced cfDNA fragments are mapped to TFBSs ±50 bp, varying numbers of short and long cfDNA fragments are found at the three TFBSs shown in (**A**). **C**) cfDNA fragment length distribution is estimated at each TFBS (purple bars) and smoothed using kernel density estimation (green line). **D**) K-means clustering is performed on smoothed length distribution to group TFBSs with similar cfDNA fragment length distribution. Here, smoothened length distributions of clusters of CTCF TFBS are shown. Weighted length (W.L.) for each CTCF length cluster is shown in parentheses.

Regular turnover of lymphoid/myeloid cells in the human body is the major contributor to the pool of cfDNA in plasma (*30*). However, in the presence of cancer, a detectable fraction of cfDNA also arises from tumors (*31, 32*). This suggests that cfDNA has the potential to map the tumor epigenome in real-time, and therefore can help uncover the regulatory landscape of cancer from plasma. Here, we map TF footprints in plasma cfDNA by combining library protocols that enrich for short fragments with computational methods that identify the subset of TFBS that leave footprints in plasma. We show that the strength of TF footprints in plasma is proportional to the binding strength of the TF in the tissue-of-origin of the cfDNA fragments, which can enable the mapping of regulatory landscapes of tumors from plasma. As proof of principle, we demonstrate that plasma TF footprints in an estrogen receptor positive (ER+) breast cancer model can predict TF-specific accessibility across human tumors, which raises the possibility of mapping tumor TF binding in human plasma. We then identify TFBS where the density of TF footprints in human plasma samples can be used to identify the presence of breast cancer. ER+ breast cancer is one of many examples of a TF driven disease: the cancer state, that is, response or resistance to drug is reflected by where in the genome ER (a TF) and related TFs like FOXA1 can bind in tumor cells (*33-35*). Thus, our results show that plasma cfDNA contains TF binding information that is specific to tumor state.

## Results

### Unique cfDNA fragment length distributions identify TF binding in the tissue-of-origin

ChIP-seq and CUT&RUN applied to cell lines and tissue samples represent gold standard methods of determining TF binding across the genome. To study human disease, it is impractical and nearly impossible to perform repeat analyses on biopsy tissues. We therefore set out to develop an alternative to ChIP-seq and CUT&RUN that can be applied to physiological and pathological states of humans in a minimally invasive manner by inferring specific TF binding from plasma cfDNA. TF footprints (<80 bp) are too short to be captured by standard library protocols, but single strand library protocol (SSP) for cfDNA can robustly capture short as well as longer, nucleosomal cfDNA fragments (*23*). In all our analyses, we used cfDNA sequencing datasets generated using SSP in this study as well as from a published study (*23*).

To ask if we can uncover TF-nucleosome dynamics from plasma cfDNA, we undertook a candidate approach of examining binding sites of specific TFs. We started with CTCF as it is constitutively expressed (*36, 37*), has a long residence time on DNA (*38*), and has known binding profiles in a large, diverse set of cell types (*18*). We aggregated CTCF binding sites from 18 cell types (70 cell lines) and analyzed fragment length distributions of cfDNA from a healthy donor (IH02 dataset (*23*)) at these sites. At each TFBS, we mapped cfDNA fragment midpoints (**Figure 1B**) and estimated a fragment length distribution (**Figure 1C**). K-means clustering of these fragment length distributions identified two types of clusters – one enriched with short cfDNA fragments (<100 bp; cluster 1 and 2) and the other enriched with long cfDNA fragments (>120 bp; cluster 3-6) (**Figure 1D**). When we mapped enrichment of cfDNA fragments around 1 kb of the TFBS, clusters 1 and 2 showed strong enrichment of short protections at TFBSs relative to 1 kb upstream and downstream of the TFBS (**Figure 2A**). Strikingly, these two clusters also showed strong nucleosome phasing at least 1 kb upstream and downstream of the TFBS (**Figure 2B**). It is well known that CTCF binding organizes nucleosomes in its vicinity (*39, 40*). Thus, fragment length profile at CTCF binding sites not only identified TF binding, but also uncovered chromatin structure surrounding the bound CTCF from plasma cfDNA. Since most cfDNA in a healthy donor arises from lymphoid/myeloid cells, we asked if the TFBS clustering based on cfDNA reflected nucleosome positioning in a representative lymphoblastoid cell line (GM12878). MNase-seq data (*18*) from GM12878 showed strong nucleosome phasing for clusters 1 and 2, but the rest of the clusters had very weak or no phasing patterns (**Figure 2C**). This strongly suggests that we can capture CTCF binding and associated nucleosome landscape from lymphoid/myeloid cells in cfDNA and that the mechanism of DNA release from these cell types gives a signal similar to MNase profiling.

**Figure 2.**
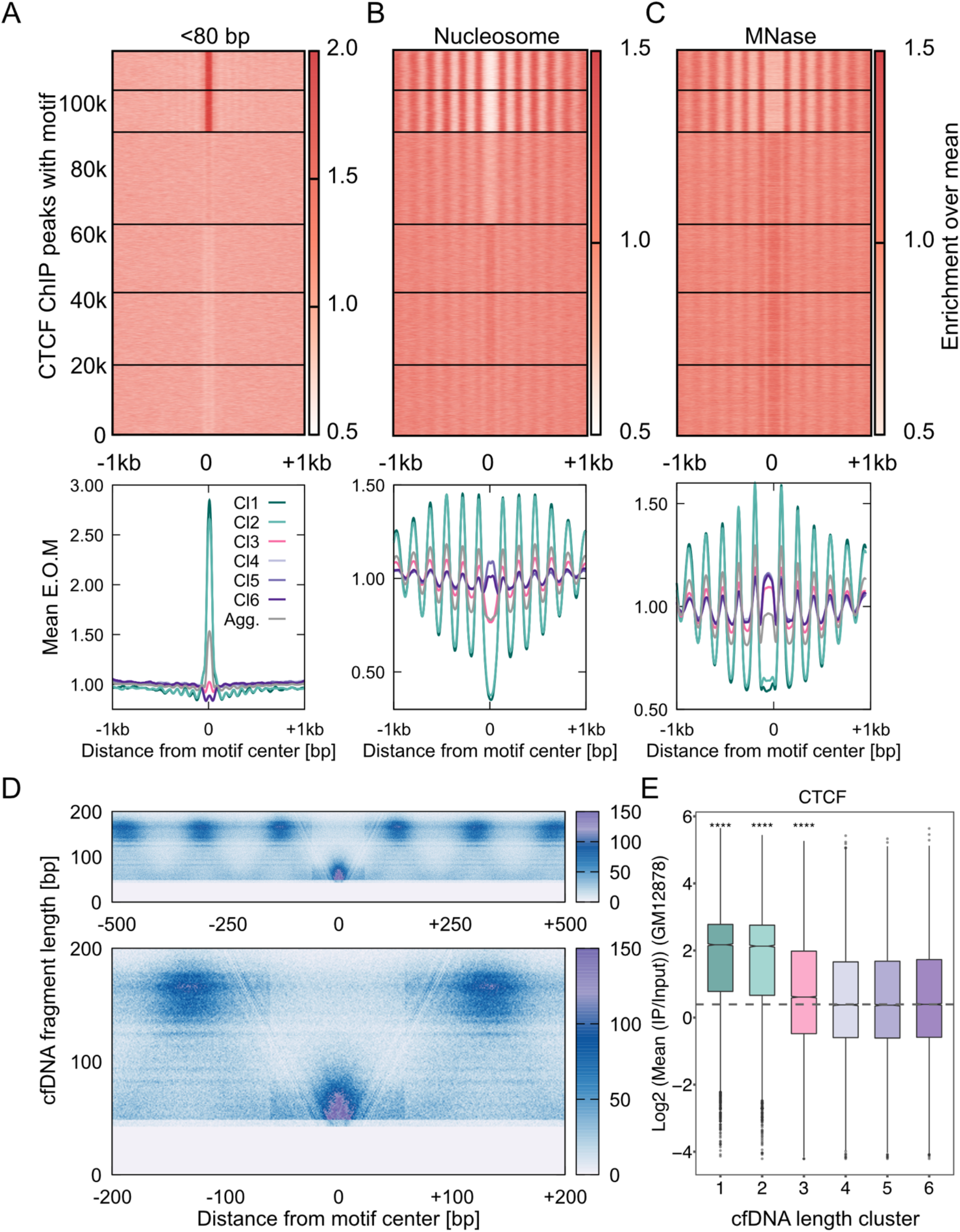
cfDNA maps CTCF-nucleosome dynamics in plasma from a healthy individual. **A**) Enrichment over the mean signal in TFBS ± 1Kb of cfDNA short (<80 bp) fragments is plotted as a heatmap (**top**, 117,144 CTCF TFBS) and as metaplots for each cluster (**bottom**). **B**) Same as (**A**) for nucleosome-sized fragments (130-180 bp). **C**) Same as (**B**) for MNase-seq dataset from GM12878 cells. Fragment midpoint versus fragment length plot (V-plot) of cfDNA fragments centered at CTCF binding sites from clusters 1 and 2. Fragment densities at motif center ± 500 bp (**top**) and motif center ± 200 bp (**bottom**) are plotted. **E**) Boxplot of CTCF mean ChIP signal from the GM12878 cell line across length clusters. Number of sites (n) in length clusters and p-value using Kolmogorov–Smirnov (KS) test with alternative = “greater” option are: Cl1: n = 11978, p (1,6) < 2.2×10^−16^; Cl2: n = 12811, p (2,6) < 2.2×10^−16^; Cl3: n = 28132, p (3,6) = 1.1×10^−31^; Cl4: n = 20839, p (4,6) = 0.95; Cl5: n = 22087, p (5,6) = 0.96; Cl6: n = 21297. p (a,b) denotes p-values calculated between scores in length cluster “a” and scores in length cluster “b”. Significance string (****) is added if p<0.0001 after Bonferroni correction.

To further visualize the chromatin structure around CTCF bound sites and identify the minimum protection conferred by CTCF on DNA, we plotted the count of cfDNA fragment midpoints around CTCF bound sites as V-plots for sites in clusters 1 and 2 (*41*). With the V-plot spanning TFBS ± 500 bp, we observe strongly positioned nucleosomes with protection length between 140-180 bp, flanking short protections at the CTCF sites in the center (**Figure 2D, top**). In the V-plot spanning TFBS ± 200 bp, a strong “V” is evident at the center, where there is an enrichment of fragments <80 bp. A “V” indicates a well-positioned, strong barrier to nucleases, which further confirms that cfDNA is directly mapping TF binding and its associated nucleosome landscapes from the cells of origin (**Figure 2D, bottom**).

The separation of bound and unbound sites by our clustering approach is also apparent when we compare the short and nucleosomal fragment enrichment at individual clusters to the aggregate enrichments across all sites (gray lines in **Figure 2A, bottom**). TF enrichment, nucleosome occlusion, and nucleosome ordering are substantially weaker in aggregate compared to clusters 1 and 2 as expected. In other words, identifying the subset of sites that are bound could inform us of TF binding strength in cfDNA cells of origin. To test this idea, we calculated the ChIP scores from GM12878 cells at TFBS belonging to each cfDNA length cluster. We found the ChIP scores of the first two clusters to be almost four times higher than the other four clusters (**Figure 2E**). The fact that hematopoietic ChIP scores correlate with our inferred sites of CTCF binding in cfDNA supports the conclusion that cfDNA length profile at TFBS reports on TF binding strength in cfDNA tissue-of-origin.

### Binding sites of hematopoietic TFs are sensitive to changes in cfDNA tissues-of-origin

Since most cfDNA in healthy individuals is of lymphoid/myeloid origin, we asked if we can map protections for lymphoid/myeloid-specific TFs: PU.1, a pioneer factor that plays a crucial role in myeloid and B-cell development (*42, 43*) and LYL1, an important factor for erythropoiesis (*44*) and development of other hematopoietic cell types (*45, 46*). Upon clustering the binding sites of PU.1 and LYL1 based on cfDNA length distributions, we found an enrichment of short protections at a subset of binding sites similar to CTCF (clusters 1 and 2; **Figure 3A, F**). Distribution of longer fragments around the binding sites showed strong nucleosomal phasing in clusters 1 and 2 (**Figure 3B, G**). The presence of nucleosome phasing further confirmed specific TF binding as this is a known outcome of LYL1 and PU.1 binding to DNA (*29, 47-49*). Clusters 1 and 2, which had the highest enrichment of short protections also had significantly higher ChIP scores in lymphoid/myeloid cells-lines compared to cluster 6 (nucleosomal) for both PU.1 and LYL1 (**Figure 3C, H**). Thus, we can map binding of hematopoietic TFs in plasma cfDNA in humans.

**Figure 3.**
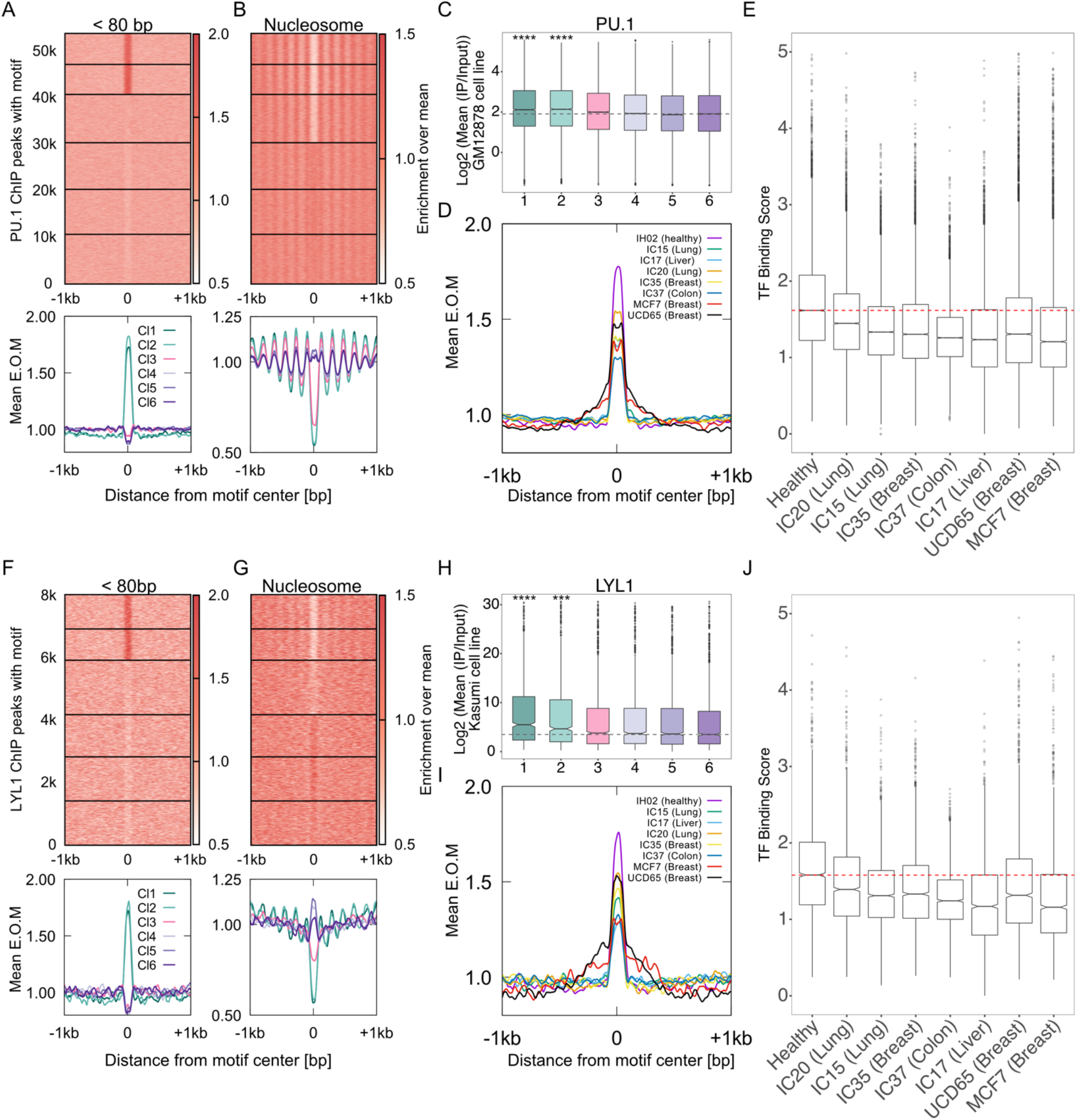
cfDNA of lymphoid/myeloid origin contains hematopoietic TF footprints. **A**) Enrichment over the mean signal in PU.1 TFBS ± 1Kb of cfDNA short (<80 bp) fragments is plotted as a heatmap (**top**, 53,613 PU.1 TFBS) and as metaplots for each cluster (**bottom**). **B**) Same as (**A**) for nucleosome-sized fragments (130-180 bp). **C**) Boxplot of PU.1 mean ChIP signal (Log2) from GM12878 cell line across length clusters. Number of sites (n) in length clusters and p-value using KS test are: Cl1: n = 6528, p (1,6) = 9.2×10^−20^; Cl2: n = 6447, p (2,6) = 1.7×10^−22^; Cl3: n = 10377, p (3,6) = 0.00011; Cl4: n = 10036, p (4,6) = 0.19; Cl5: n = 9673, p (5,6) = 0.7; Cl6: n = 10552. Significant string was determined after Bonferroni correction. **D**) Enrichment metaplots for short fragments in PU.1 TFBS belonging to clusters 1 and 2 for healthy (IH02), cancer (IC15, 17, 20, 35, and 37) cfDNA and PDX cfDNA (MCF7 and UCD65). Boxplot of mean of short fragment enrichment (TFBS±50 bp) for the samples and TFBS plotted in (**D**).Same as (**A**) for LYL1 (7,999 TFBS). **G**) Same as (**B**) for LYL1. **H**) Same as (**C**) for LYL1. Number of sites (n) in length clusters and p-value using KS test are: Cl1: n = 1083, p (1,6) = 4.7×10^−12^; Cl2: n = 1001, p (2,6) = 3×10^−7^; Cl3: n = 1748, p (3,6) = 0.18; Cl4: n = 1351, p (4,6) = 0.15; Cl5: n = 1415, p (5,6) = 0.62; Cl6: n = 1401. Significant string was determined after Bonferroni correction. **I**) Same as (**D**) for LYL1. **J**) Same as (**E**) for LYL1. ****: p<0.0001, ***: 0.0001<p<0.001

In cancer patients, cancer cells also contribute significantly to plasma cfDNA. Hence, we hypothesized that cancer cell derived cfDNA will lead to dilution of lymphoid/myeloid signal. Such dilution would lead to a proportional decrease in enrichment of short fragments at Clusters 1 and 2 of hematopoietic TFBS due to cfDNA contributions from non-hematopoietic cell types where PU.1 and LYL1 are absent. To test this hypothesis, we performed k-means clustering of PU.1 and LYL1 binding sites based on the cfDNA length distributions for cfDNA from donors with cancer. We found that the short fragment enrichment for the bound clusters (1 and 2) was the highest for healthy human plasma (**Figure 3D, E, I, and J**). Cancer samples had significantly weaker short fragment enrichment at sites from clusters 1 and 2 for PU.1 and LYL1 (**Figure 3E, J**) and did not have higher ChIP scores compared to cluster 6 (**Supplementary Figure 1A-D**). In addition to using cfDNA from cancer patients, we also used human cfDNA from cell-line/patient-derived xenografts (C/PDXs) (**Figure 4A**). Since the only source of human cfDNA in a xenograft is from the cancer cells, fragments that uniquely map to the human genome in this context represent pure circulating tumor DNA (ctDNA). We found no expression of PU.1 or LYL1 in breast tumor model systems, and accordingly, we observed no nucleosome phasing or higher ChIP scores for the top 2 clusters in the xenograft cfDNA (**Supplementary Figure 1E-H**). Additionally, we found an expected decrease in enrichment of short fragments in clusters 1 and 2 from the xenografts when compared to healthy donor (**Figure 3D, I, E, and J;** sample names: UCD65 and MCF7). The clear separation between cfDNA from a healthy donor and cfDNA from cancer patients and from xenografts suggest that the length profiles of cfDNA at hematopoietic TFBS when combined with local enrichment of short fragments can identify dilution of lymphoid/myeloid cfDNA across diverse plasma samples.

**Figure 4.**
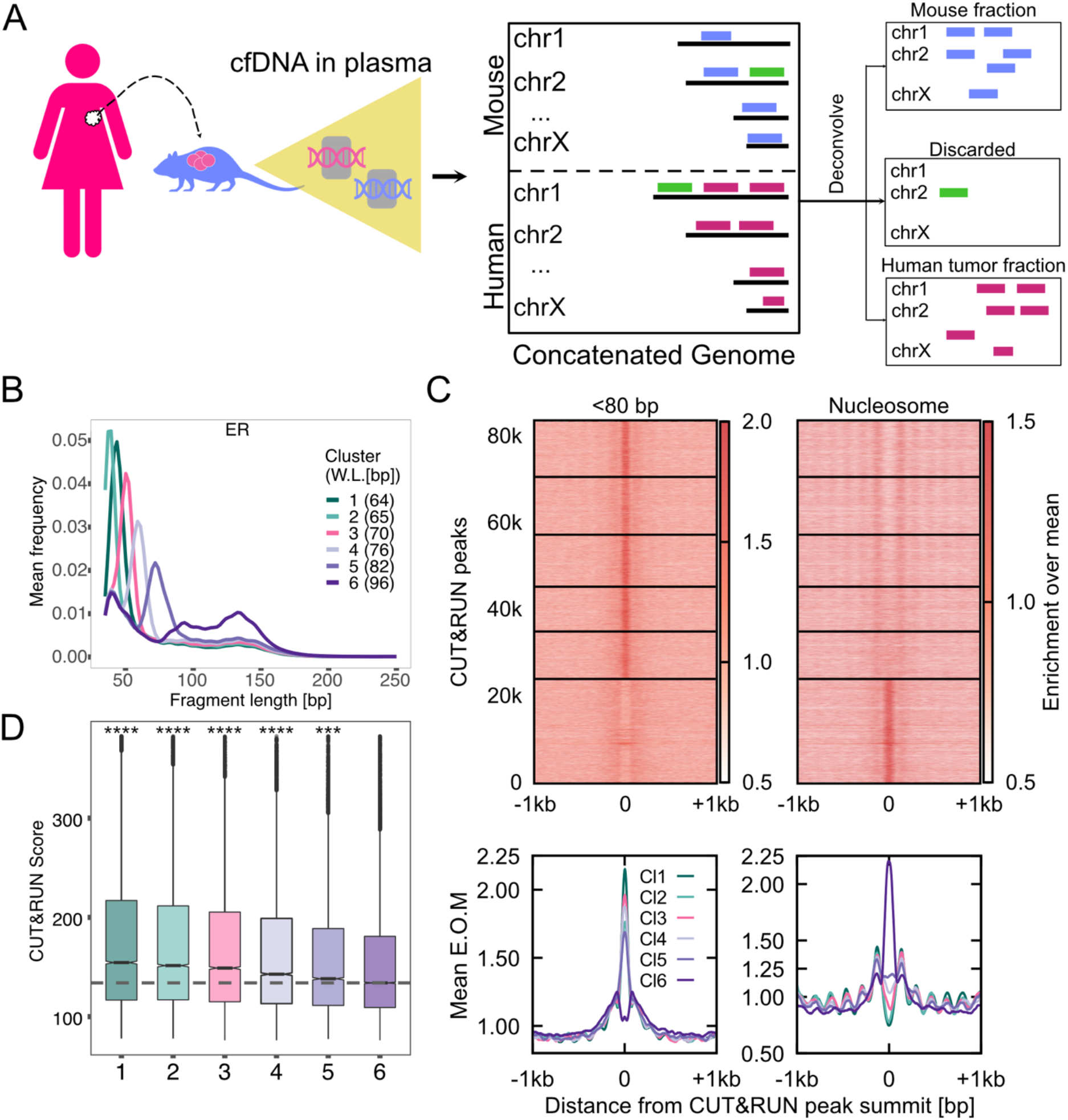
ER+ PDX models enable identification of pure tumor cfDNA footprints for ER. **A**) Schematic of human tumor implant in mouse and the process of identifying tumor cfDNA by mapping mouse plasma cfDNA to an *in silico* concatenated genome. Fragments mapping uniquely to human (violet lines) defines tumor cfDNA (ctDNA). Fragments mapping uniquely to mouse genome (blue lines) arise from the tumor microenvironment and from the mouse lymphoid/myeloid cells. Fragments mapping to both genomes were discarded (green lines). **B**) Average length distributions at clusters of ER CUT&RUN peaks (summit ± 50 bp) generated by k-means clustering (n = 6) of the ctDNA fragment length distribution. **C**) Enrichment over the mean signal in TFBS ± 1Kb of cfDNA short (<80 bp) fragments is plotted as a heatmap (**top**, 83,311 ER TFBS) and as metaplots for each cluster (**bottom**). **D**) Boxplot of ER CUT&RUN scores for peak summits in k-means clusters. Number of sites (n) in length clusters and p-value using KS test are: Cl1: n = 12785, p (1,6) = 1.2×10^−151^; Cl2: n = 13301, p (2,6) = 7.9×10^−116^; Cl3: n = 11943, p (3,6) = 1.5×10^−80^; Cl4: n = 10363, p (4,6) = 1.6×10^−37^; Cl5: n = 10848, p (5,6) = 1.1×10^−08^; Cl6: n = 24029. Significant string was determined after Bonferroni correction. ****: p<0.0001, ***: 0.0001<p<0.001

### ctDNA maps tumor-specific TF binding

We were able to uncover strong signals of CTCF and hematopoietic TFs binding in plasma cfDNA because the vast majority of cells that release cfDNA have these TFs bound in their genome. However, tumor-specific TFs will, by definition, have weaker signals because tumor cfDNA is always a minor fraction of total cfDNA. In order to develop pure tumor signatures of TF binding in cfDNA, we turned to human cancer xenografts implanted in mice. Since the tumor-derived cfDNA in PDX would map to the human genome, and the endogenous cfDNA from the mouse would map to the mouse genome, we could identify cfDNA molecules from sequencing that were purely from the tumor, hence circulating tumor DNA (ctDNA), but obtained from a closed *in vivo* system (**Figure 4A**). We used ER+ breast tumor cells, UCD65 (*50*) and MCF7, as ER+ tumors are driven by the TFs Estrogen Receptor (ER) and FOXA1. We first profiled ER and FOXA1 binding using CUT&RUN (*19*). CUT&RUN is an alternative to ChIP-seq that relies on a protein-A-tagged nuclease that binds to a primary antibody of epitope of choice. The nuclease is activated upon addition of calcium, which results in the release of DNA fragments bound to ER. Due to the absence of crosslinking and release of bound sites rather than enrichment of bound sites, CUT&RUN captures TF binding at higher sensitivity and provides a greater dynamic range of signals compared to ChIP-seq (*19*). We performed CUT&RUN for ER and FOXA1 in estradiol (E2)-treated MCF7 cells and obtained ∼80,000 and ∼40,000 CUT&RUN sites for ER, and FOXA1 respectively, with sufficient coverage in our PDX cfDNA datasets (MCF7, UCD65).

Importantly, when we performed fragment-length distribution analysis at ER CUT&RUN peaks and defined six clusters, the four clusters with lowest expected fragment length (**Figure 4B**) showed strong short fragment protections and phased nucleosomes (**Figure 4C**) as well as significantly higher ER binding measured as CUT&RUN score (**Figure 4D**). We observed similar trends for FOXA1 binding sites (**Figure 5A-C**). Positive correlation between ctDNA short fragment enrichment and CUT&RUN scores strongly suggests that we are capturing binding in cancer cells and that the signal from cfDNA release *in vivo* is similar to CUT&RUN profiling. Thus, defining binding sites in tumor cells using CUT&RUN enables sensitive mapping in plasma of the TF-binding that occurs in tumor-cells-of-origin.

**Figure 5.**
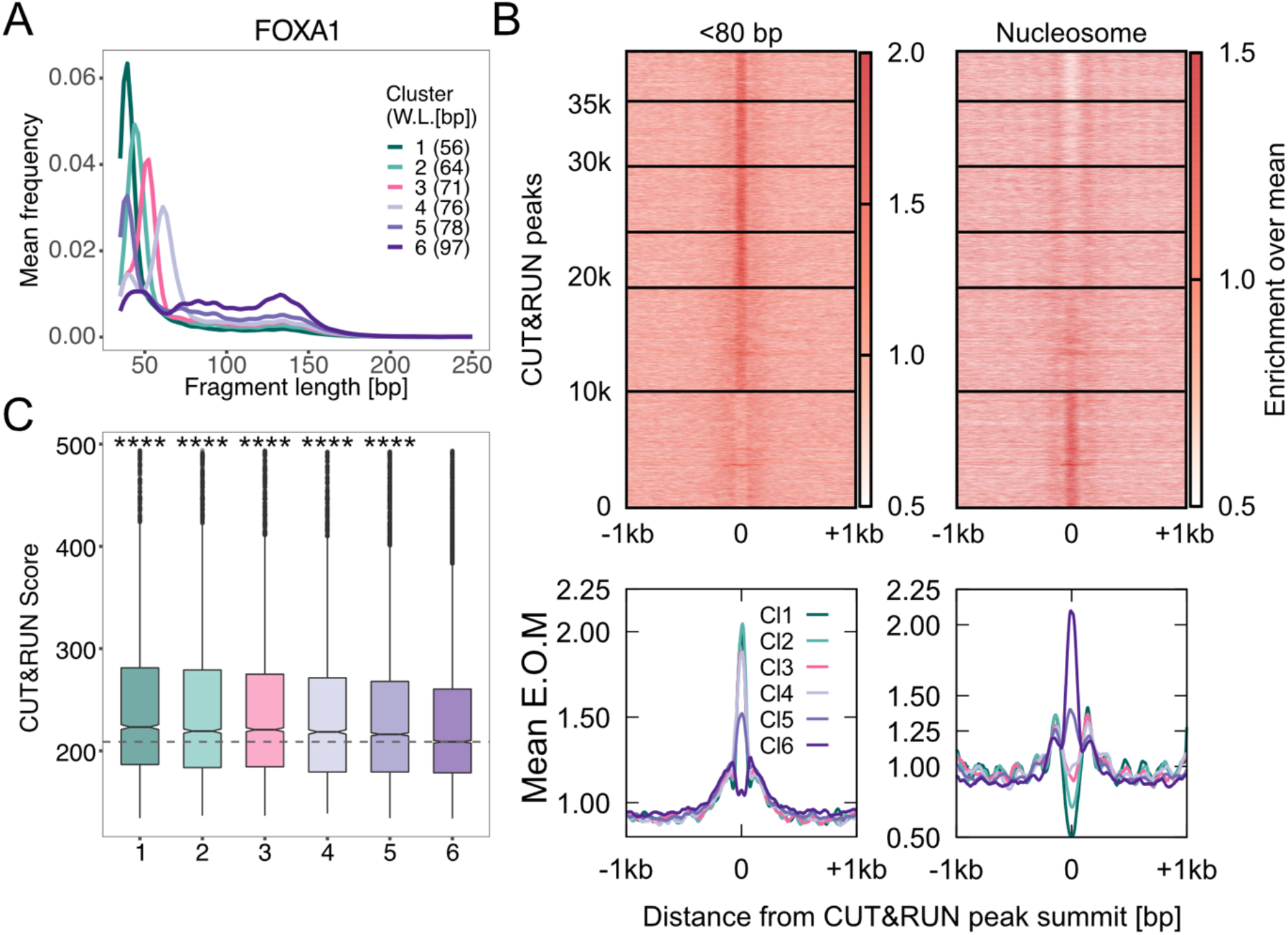
ER+ PDX models enable identification of pure tumor cfDNA footprints for FOXA1. **A**) Average length distributions at clusters of FOXA1 CUT&RUN peaks (summit ± 50 bp) generated by k-means clustering (n = 6) of the ctDNA fragment length distribution. **C**) Enrichment over the mean signal in TFBS ± 1Kb of cfDNA short (<80 bp) fragments is plotted as a heatmap (**top**, 39,500 FOXA1 TFBS) and as metaplots for each cluster (**bottom**). **D**) Boxplot of FOXA1 CUT&RUN scores (see methods) for peak summits in K-means clusters. p values from Kolmogorov–Smirnov test. Number of sites (n) in length clusters and p-value using KS test are: Cl1: n = 4220, p (1,6) = 3.4×10^−36^; Cl2: n = 5669, p (2,6) = 3.2×10^− 19^; Cl3: n = 5699, p (3,6) = 4.5×10^−15^; Cl4: n = 4831, p (4,6) = 3.1×10^−10^; Cl5: n = 9033, p (5,6) = 3.9×10^−10^; Cl6: n = 10017. Significant string was determined after Bonferroni correction. ****: p<0.0001.

### Unique sets of TFBS display tissue-of-origin-specific TF protections in plasma

We have defined sets of binding sites that show TF-specific protections in two pure systems: healthy plasma and PDX plasma. We now asked if we could define subset of these sites that would be unique to the tissue-of-origin. To do this, we performed length clustering analysis at all TFBS with both healthy plasma dataset and with the PDX datasets to identify binding site clusters with significantly higher ChIP/CUT&RUN binding scores compared to the nucleosomal cluster of binding sites for each cfDNA dataset. We then intersected the significant binding sites between healthy plasma and PDX models. First, we found that PU.1 and LYL1 sites had TF protections that correlated with binding strength only in healthy plasma (**Figure 6A**), indicating that all significant TFBS of PU.1 and LYL1 could be used to identify hematopoietic contribution to cfDNA. CTCF is a constitutive factor, ER is expressed in T cells (*51, 52*), and factors related to FOXA1 that have same binding motifs are expressed in hematopoietic cells, for example, FOXM1 (*53-55*). The partial overlap of binding of these or related factors in hematopoietic and cancer cells led to us finding sites with significant TF protections in both healthy plasma and in PDX for CTCF, FOXA1, and ER (**Figure 6A, Supplementary Figures 2, 3**). For example, a large fraction of sites of CTCF (16709 in set 2 and 4945 in set 4) are shared between PDX and healthy plasma. Rest of the CTCF sites (17902 in set 1, 6022 in set 3, 4930 in set 5, and 4649 in set 6, CTCF in **Figure 6A**) are cancer specific. In contrast, the top 3 sets of sites for FOXA1 and ER are PDX-specific, with the largest set of sites specific to UCD65 (8226 for FOXA1 and 13879 for ER). FOXA1 has sites specific to MCF7 as well (set 3) and ER has sites specific to MCF7 (set 3) and UCD4 (set 6). Thus, in spite of overlap in binding between hematopoietic cells and cancer cells, ER and FOXA1 have enough unique sites protected in plasma that not only distinguish healthy plasma from PDX, but also distinguish individual PDX’s.

**Figure 6.**
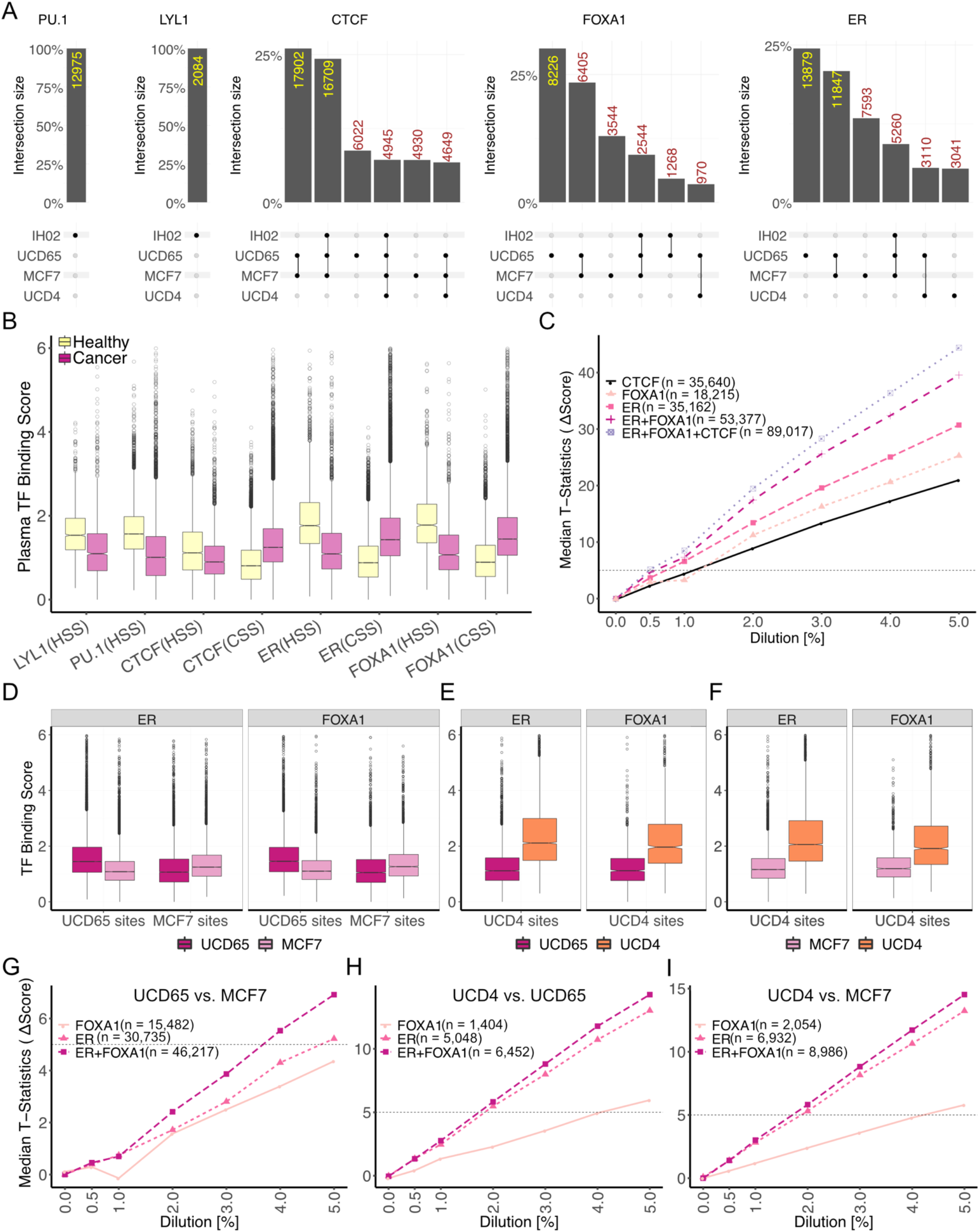
Tissue-specific TF binding sites enable detection of disease states. **A)** Upset plots (*75*) of cfDNA-inferred bound sites in different plasma samples for LYL1, PU.1, CTCF, FOXA1 and ER (left to right). Plots were generated using ComplexUpset R package (DOI: 10.5281/zenodo.4661589). **B)** Boxplots of TF binding scores measured as mean enrichment of short fragments at CUT&RUN peak summit ± 100 bp for ER and FOXA1 and motif center ± 50 bp for LYL1, PU.1 and CTCF. CSS – cancer specific sites; HSS; healthy specific sites **C)** Line plot of median t-statistic calculated for change in the binding scores (score in healthy plasma used as baseline) at binding sites of an individual TF or a collection of TFs at different *in silico* dilutions of healthy cfDNA with PDX ctDNA. At each dilution, 100 bootstrapped samples were generated. Horizontal dashed line is drawn where the t-statistic equals 5. **D)** Boxplot of TF binding scores in pure ctDNA (UCD65/MCF7) at ER and FOXA1 sites specific to UCD65 or MCF7. **E)** Boxplot of TF binding scores in pure ctDNA (UCD65/UCD4) at ER and FOXA1 sites specific to UCD4 against UCD65. **F)** Boxplot of TF binding scores in pure ctDNA (MCF7/UCD4) at ER and FOXA1 sites specific to UCD4 against MCF7. **G)** Line plot of median t-statistic calculated for the change in TF binding scores at UCD65 or MCF7-specific ER, FOXA1, or for ER and FOXA1 sites combined. **H)** Same as (**G**) for UCD4-specific ER and FOXA1 sites against UCD65. **I)** same as (**G**) for UCD4-specific ER and FOXA1 sites against MCF7.

Although FOXA1 is not expressed in lymphoid/myeloid cells, some FOXA1 binding sites identified in MCF7 cells showed significant enrichment of TF footprints in healthy plasma. We asked if related FOX factors like FOXM1 and FOXK2 that are expressed in lymphoid/myeloid cells may be binding at these sites to give rise to short footprints in cfDNA. To ask if FOXM1 or FOXK2 give rise to footprints at a subset of FOXA1 sites, we calculated scores for FOXM1 and FOXK2 binding from ChIP experiments conducted in GM12878 cells. We found FOXM1 ChIP scores to strongly correlate with short length clusters in healthy plasma but not FOXK2 ChIP scores (**Supplementary Figure 3**). This suggests that FOXM1 occupies sites in lymphoid/myeloid cells that are a subset of sites bound by FOXA1 in MCF7 cells.

With these collections of sites that were unique to cancer and to the ER status (normal vs. amplified vs. mutated), we calculated a plasma TF binding score: the number of short reads (<80 bp) mapped within 50 bp of the TFBS normalized by the number of reads in 1000 bp around the TFBS. This plasma TF score tracks with the identity of the sites: the sites unique to healthy plasma had a significantly higher TF score for healthy plasma compared to PDX and vice versa. Similarly, sites specific to UCD65, MCF7, and UCD4 when compared to each other also had higher plasma TF scores (**Figure 6B, D, E, and F**). Thus, unique sets of sites identified using cfDNA length clusters also had localized enrichment of short fragments relative to the surrounding 1000 bp in a system-specific manner, which shows the potential of cfDNA length clusters to identify not only the tissue-of-origin but also the disease state.

In a plasma sample from an individual with cancer, both lymphoid/myeloid cells and tumor cells will contribute to cfDNA, with majority of the contribution still being from the lymphoid/myeloid cells. To ask at what dilution of tumor DNA we could detect the presence of cancer using TF footprints, we performed *in silico* dilutions of PDX cfDNA, which represents pure tumor DNA into healthy plasma cfDNA at 0, 0.5, 1, 2, 3, 4, and 5%. We then calculated plasma TF binding score at sites specific to healthy plasma and PDX. We compared these scores between the *in silico* diluted plasma samples and non-diluted plasma sample to calculate a paired t-statistic. We set a cut-off of 5 for the median paired t-statistic to indicate a significant difference between diluted and non-diluted plasma sample (**Supplementary Figure 4**). We found ER sites to be strongest in separating tumor diluted cfDNA from pure healthy cfDNA (detection at <1% tumor cfDNA) followed by FOXA1 and CTCF (detection at ∼1% of tumor cfDNA, **Figure 6C**). PU.1 (detection at 2% tumor cfDNA) and LYL1 had weaker but significant contributions (**Supplementary Figure 5**). Combined ER and FOXA1 sites showed a median t-statistic greater than 5 between 0.5 and 1% tumor fraction. Since most metastatic disease states have tumor fractions higher than 1% (*56, 57*), our analysis suggests that we would be able to delineate TF binding in metastatic tumors, in spite of the significant interference from cfDNA of lymphoid/myeloid origin.

We next asked if we could differentiate between the PDXs based on their ER status: ER expression is much higher in UCD65 (ESR1 amplification) and UCD4 has a mutated ER (activating D538G mutation) (*58*). Both ER and FOXA1 sites contribute to differentiating UCD65 from MCF7. Combining sites from both TFs is synergistic and separates UCD65 and MCF7 at 4% of tumor fraction (t-statistic > 5, **Figure 6G**). Thus, at marginally higher tumor fractions, we can even identify signatures of differences in ER expression levels using TFBS defined by a combination of CUT&RUN and cfDNA length clustering. Strikingly, ER sites could robustly differentiate UCD4 from UCD65 and MCF7 (**Figure 6H, I**), highlighting the fact that mutated ER leads to differential binding signature that can be identified in plasma cfDNA at 2% tumor fraction. Significantly, FOXA1 sites were much weaker than ER in differentiating UCD4 from UCD65 and MCF7, highlighting that the mutation-specific changes in TF footprints in plasma is strongest for ER. In summary, by identifying the subset of high-resolution TFBS protected in distinct plasma samples, we are able to define TF signatures unique to ER+ breast cancer and further, unique to amplified WT ER and ER D538G.

### Identified TFBS report on tumor TF binding in individuals with breast cancer

Since our *in silico* dilution analyses indicate that TF footprints in plasma can identify breast cancer disease state at tumor fractions of 1-4%, we next asked if the TFBSs we identified to be uniquely protected in PDX plasma would reflect disease states in heterogeneous human samples. To test this, we first turned to ATAC-seq datasets generated using primary tumor samples in the TCGA database. ATAC-seq reports on DNA accessibility, which highly correlates with TF binding (*59*). We asked if tumors exhibited TF-specific accessibility at the TFBSs we identified. We ordered BRCA tumors based on a specific TF’s expression and then calculated accessibility at sites identified to be UCD65-specific. We found tumors that express ER (Transcripts Per Million (TPM) ≥ 10) had a vast majority of UCD65-specific ER sites with higher accessibility compared too tumors that do not express ER (TPM < 10, **Figure 7A**). We found even stronger accessibility differences at UCD65-specific FOXA1 binding sites, with FOXA1-expressing tumors having much higher ATAC scores than FOXA1-non-expressing tumors at a vast majority of sites (**Figure 7B**).

**Figure 7.**
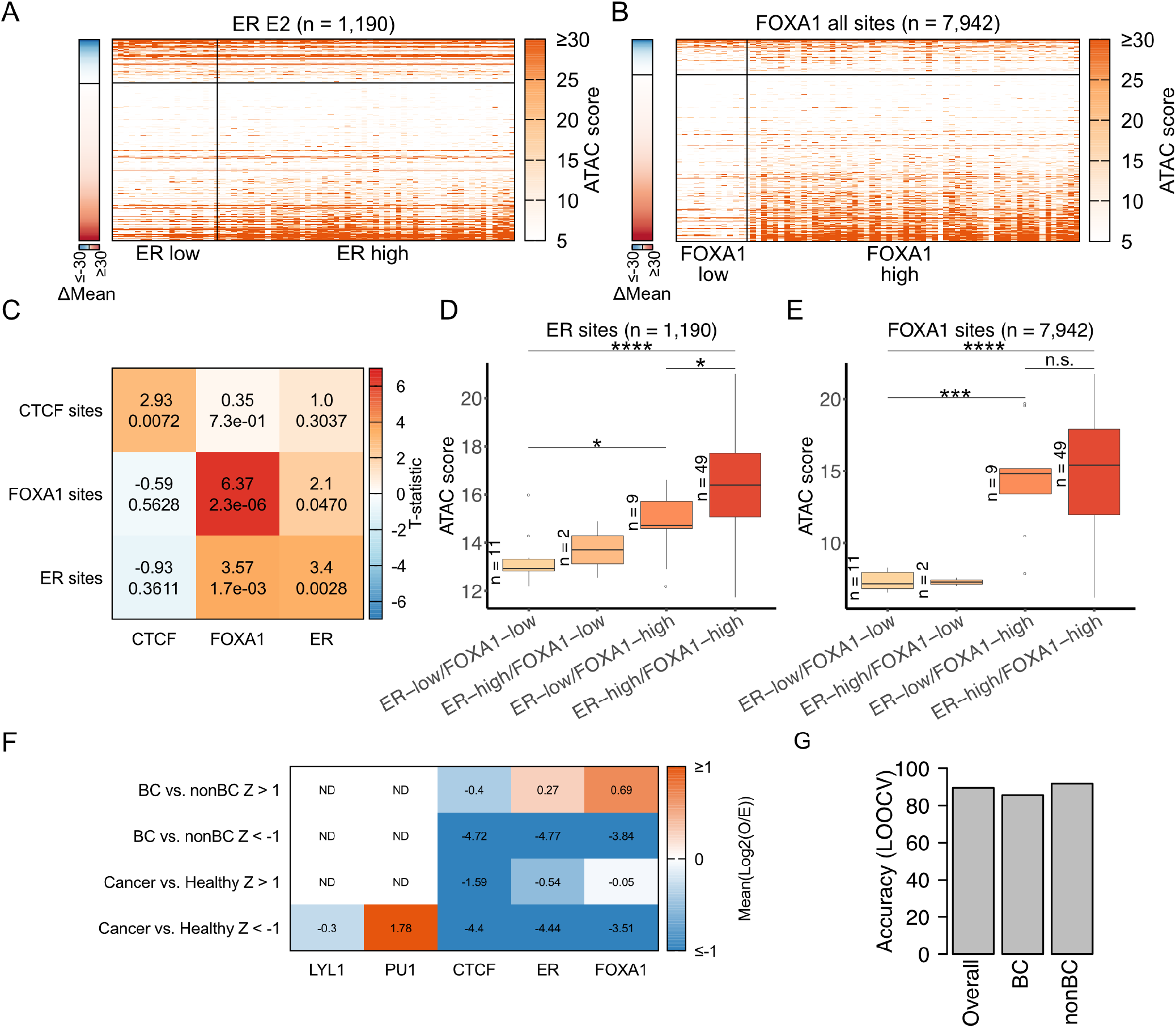
Plasma footprints represent TF specific accessibility in primary tumors and can predict presence of breast cancer. **A**) Heatmap of ATAC scores from BRCA cohorts stratified based on ER expression levels (ER low: TPM < 10, ER high: TPM ≥ 10) at cfDNA-inferred ER CUT&RUN peaks with ER motif. The single column heatmap (**left**) plots the difference in mean ATAC scores between tumors with high ER expression and tumors with and low ER expression. The sites are ordered in ascending order of difference in ATAC scores between the two groups and the horizontal line separates sites with higher score in ER high compared to ER low. **B**) Same as (**A**) for FOXA1 sites. **C**) Heatmap of t-statistic calculated between tumors grouped by TF expression (columns; low (bottom 15 cohorts) and high (top 15 cohorts) expression levels) at binding sites of different TFs (rows). **D**) Boxplot of mean ATAC-scores at ER sites (n = 1,190) where tumors are stratified by both ER and FOXA1 expression. **E**) Boxplot of mean ATAC-scores at FOXA1 sites (n = 7,942) where patients are stratified by both ER and FOXA1 expression. **F**) Heatmap of enrichment (Log2 (Observed/Expected)) of frequency of TF features selected for a given classification (rows) divided by overall frequency of TF features. **G**) Prediction accuracy of classifying patients to BC (breast cancer) and nonBC (non-breast cancer) using TF scores from plasma cfDNA using leave one out cross-validation.

FOXA1 is known to act as a pioneer factor, enabling ER binding by establishing accessibility at its binding sites (*34, 60*). We asked if we could reproduce this finding at ER and FOXA1 binding sites we identified by taking advantage of the heterogeneity in ER and FOXA1 expression across TCGA samples. If the ER and FOXA1 sites we identified are representative of ER and FOXA1 function across human breast tumors, then accessibility at ER binding sites should depend on the presence of FOXA1. CTCF is a good control as its expression shouldn’t influence accessibility at ER or FOXA1 sites. We first calculated the mean ATAC-score for each tumor sample by aggregating the ATAC score across all sites of a given TF. For CTCF, ER, and FOXA1 sites, we performed two sample t-test (sample 1: cohorts with high TF expression (top 15), sample 2: cohorts with low TF expression (bottom 15)). We found the mean ATAC-scores at CTCF, FOXA1, and ER sites were significantly different when tumors were grouped by the expression of the respective TF, with strongest difference seen for FOXA1 (diagonal cells in **Figure 7C**). Strikingly, we observed a strong difference (t-statistic = 3.57; p = 1.7×10^−3^) in mean ATAC-scores at ER sites when tumors were grouped based on FOXA1 expression. This difference was stronger than at FOXA1 sites when tumors were grouped based on ER expression (t-statistic = 2.1; p = 0.047), suggesting that FOXA1 expression has a stronger influence on accessibility at ER sites than *vice versa*.

To further explore the effect of FOXA1 at ER sites, we stratified BRCA tumors by both ER and FOXA1 expression levels. In tumors with low ER expression, increase in FOXA1 expression led to a significant increase in mean ATAC-scores at ER sites, suggesting that FOXA1 keeps the chromatin open at ER sites even in the absence of ER (**Figure 7D**). Expression of ER and FOXA1 led to the highest accessibility at ER sites suggesting further chromatin opening post ER binding (**Figure 7D**). In stark contrast, at FOXA1 sites, accessibility increase is seen only due to increase in FOXA1 expression. Thee presence of ER did not lead to a significant increase in accessibility (**Figure 7E**). Our observation of FOXA1 expression driving accessibility at both ER and FOXA1 binding sites agrees well with the fact that FOXA1 is a pioneer factor that opens up ER sites. Taken together, our analysis shows that sites with tumor-specific plasma protections in PDXs can define TF-specific accessibility across human breast tumors. These results indicate that TF protections in plasma can define tumor TF binding in humans.

Next, we asked if TF binding scores from plasma cfDNA can distinguish cancer from healthy states and breast cancer from other cancers and healthy states. We compared TF binding scores in 19 human plasma cfDNA sequencing datasets (healthy = 4, non-breast cancer = 8 (total nonBC = 12); breast cancer (BC) = 7). To take advantage of samples that were sequenced at varying depths, we defined TF features as aggregates of 250 binding sites of the TF after ordering all its binding sites by ChIP/CUT&RUN score. We ended up with a total of 359 features (PU.1 = 43, LYL1 = 7, CTCF = 120, ER = 124, FOXA1 = 65). We made two classification groups: cancer vs. healthy (n=15,4) and BC vs. nonBC (n=7,12). We calculated the Z-score for each feature for these two groups of classification. We then filtered for those features with |Z| >1 in each of the two classifications as features that differentiated the two classes in each classification. We then asked which of the TFs had their features over-represented or under-represented in each classification. We found PU.1 features to be over-represented in having higher TF binding scores in healthy samples compared to cancer samples (**Figure 7F**). In classifying BC and nonBC, we found no TFs to be overrepresented in features that had higher binding scores in nonBC. However, ER and FOXA1 features were overrepresented with higher binding scores in BC compared to nonBC (**Figure 7F**). The fact that FOXA1 and ER binding sites can separate BC from nonBC indicates that the sites identified from PDXs are transferrable to human samples. Furthermore, in spite of dilution by cfDNA from lymphoid and myeloid cells, cancer-specific TF protections in plasma are sensitive markers of disease presence. To ask how accurate these features are in identifying presence of breast cancer, we resorted to leave-one-out cross validation. We identified features that significantly separated BC from nonBC using all but one of the samples (18 out of 19) and then used these features to predict status of the left-out sample. We observed an overall prediction accuracy of 89.5%, prediction accuracy of 85.7% for BC (6/7 predicted correctly), and accuracy of 91.7% for nonBC (11/12 predicted correctly, **Figure 7G**). Thus, our analysis with low to intermediate depth sequencing of 19 human plasma samples shows potential for plasma TF footprints to identify breast cancer tissue-of-origin.

## Discussion

Interaction of TFs with DNA is fundamental to gene regulation and distinct cell types are defined by unique TF-binding profiles. It is known that cfDNA fragments in plasma maintain information regarding chromatin dynamics and TF binding (*61*). Previous approaches have averaged the coverage of short and long cfDNA fragments from 1000s of sites in order to infer binding of a single TF (*23, 29*). Such analyses on aggregated sites can be used to build diagnostic classifiers but lack the granularity to generate binding profiles of TFs specific to the cells releasing cfDNA. We hypothesized that if TF protections in tissue-of-origin lead to short footprints in plasma, then mapping these footprints at individual sites would allow us to study the TF’s function in cfDNA tissues-of-origin. To this end, we first defined the subset of all possible sites of a TF that give rise to short footprints in cfDNA. Surprisingly, we found that enrichment of short footprints in plasma for a TF binding site correlates with strength with which the TF binds that site in the cells from which the cfDNA originated. We observed this for both constitutive factors with long residence times like CTCF and for tumor-specific, dynamic factors with short residence times, such as ER. This finding elevates cfDNA from a mere classifier to a means to understand TF binding in living mammals in a minimally invasive manner and in real time.

TFs in the same family use overlapping binding sites in different cell types. For example, FOXA1, which is active in hormone-dependent cancers of the breast and prostate, shares binding motif with other FOX factors like FOXM1, whose expression is enhanced in lymphoid tissues (*54, 55*), and FOXK2, which is expressed in many tissues including lymphoid/myeloid (*62*). We show that a subset of FOXA1 sites bound in ER+ tumors also give rise to short cfDNA footprints in healthy plasma. We can predict these protections to arise from FOXM1 rather than FOXK2 binding based on correlation of the enrichment of cfDNA TF footprints with binding strengths of FOXM1 in a lymphoid cell line, underscoring our ability to uncover specific TF binding profiles directly from plasma. Similarly, ER, which drives ER+ breast cancers is also active in T cells (*51*), with shared and unique binding sites between tumors and T cells. Our inference of TF footprints at each binding site of ER from PDX and healthy plasma has enabled identification of sites in an ER+ tumor also bound by ER in lymphoid/myeloid cells. This careful characterization of ER and FOXA1 has enabled us to identify binding sites of FOXA1 and ER that are not only specific to ER+ breast cancer, but which can also distinguish basal levels of ER from ER overexpression and the presence of mutated ER. Starting only with a reference set of CUT&RUN sites from E2 treated MCF7 cells, we were able to identify ER and FOXA1 binding sites that defined lymphoid/myeloid signatures and ER+ tumor subtypes. Thus, further high-resolution characterization of TFs in the future can only improve our ability to generate binding profiles from plasma in health and disease.

Our results uncover two aspects of cfDNA biology that can vastly expand the information we can gain from its study. First, enrichment of short fragments enables accurate identification of TF footprints at single binding sites in plasma. Second, careful consideration of which sites in the genome to look at can yield cancer sensitive signatures, which enable tissue-of-origin mapping of TF binding profiles. Here, we have shown that analysis of CUT&RUN data on tumor cells combined with cfDNA data from PDX models can identify tumor TF sites that are bound in a TF-specific manner across human breast tumors. In the future, putative tumor state-specific TF sites where we can analyze cfDNA footprints could be identified by mining TCGA ATAC-seq datasets and *de novo* identification of TF binding sites from models of pure tumor cfDNA. Thus, we believe our characterization of ER+ breast tumors using cfDNA TF footprints represents the tip of the iceberg for characterizing tumor phenotypes from plasma and is applicable across disease states.

## Acknowledgements

This work was supported by the RNA Bioscience Initiative, University of Colorado School of Medicine, ACS IRG #16-184-56 from the American Cancer Society (to S. Ramachandran), Earlier.org -Friends for an Earlier Breast Cancer Test (to S. Ramachandran), National Cancer Institute grants R01CA140985 (C.A.S.), R01CA205044 (P.K.), the Breast Cancer Research Foundation 16-072 (C.A.S.), the University of Colorado Cancer Center’s Oncology Research Information Exchange Network, University of Colorado Cancer Center Pathology Shared Resource (This resource is supported by the Cancer Center Support Grant (P30CA046934)), and Colorado Lung Cancer Specialized Program of Research Excellence (P50 CA058187). S. Ramachandran is a Pew-Stewart Scholar for Cancer Research, supported by the Pew Charitable Trusts and the Alexander and Margaret Stewart Trust. A.Z. was supported by an American Cancer Society -Virginia Cochary Award for Excellence in Breast Cancer Research Postdoctoral Fellowship, PF-20-095-01-DMC. The results shown here are in part based upon data generated by the TCGA Research Network: https://www.cancer.gov/tcga. We thank Dr. Olivia Rissland and Dr. Sujatha Jagannathan for critical comments on the manuscript.

## Materials

### Plasma samples

Plasma sample information is described in **Supplementary Table 1**.

### ChIP-seq peaks

We collected ChIP-peaks from publicly available datasets (*18, 63, 64*). We obtained clustered peaks for CTCF and PU.1 from ENCODE (http://hgdownload.cse.ucsc.edu/goldenPath/hg19/encodeDCC/wgEncodeRegTfbsClustered/wgEncodeRegTfbsClusteredV3.bed.gz). For LYL1, we used peaks from ReMap (http://remap.univ-amu.fr/storage/remap2020/hg38/MACS2/TF/LYL1/remap2020_LYL1_all_macs2_hg38_v1_0.bed.gz).

### TF motifs

We used TF motifs from JASPAR (*65*) (CTCF: http://jaspar.genereg.net/matrix/MA0139.1/, PU.1: http://jaspar.genereg.net/matrix/MA0080.5, ER: http://jaspar.genereg.net/matrix/MA0112.1; http://jaspar.genereg.net/matrix/MA0112.2; http://jaspar.genereg.net/matrix/MA0112.3, FOXA1: http://jaspar.genereg.net/matrix/MA0148.1; http://jaspar.genereg.net/matrix/MA0148.2; http://jaspar.genereg.net/matrix/MA0148.3) and HOCOMOCO (*66*) (LYL1: http://hocomoco.autosome.ru/motif/LYL1_HUMAN.H11MO.0.A).

### Genome-wide signal

We used publicly available genome-wide signal files in bigwig format to map ChIP and MNase signal to TF binding sites and their flanks. CTCF: https://www.encodeproject.org/files/ENCFF578TBN/@@download/ENCFF578TBN.bigWig, PU.1: https://www.encodeproject.org/files/ENCFF324NQZ/@@download/ENCFF324NQZ.bigWig, LYL1: GEO: GSE63484.

## Methods

### cfDNA extraction

1-4 mL human plasma or 0.2-0.5 mL of mouse serum were thawed from -80°C storage. Plasma or serum were spun at max speed (21,000 rcf) at 4°C for 5-10 mins to pellet any cell debris. Supernatant was transferred to new tubes and cfDNA was extracted using the QIAGEN ccfMinElute kit (cat. 55204) and eluted in 30 µL of nuclease-free water and directly added to the single-stranded DNA library protocol (SSP) or stored at -20°C.

### Single-Stranded DNA Library Protocol (SSP)

The capture of cfDNA fragments from plasma or serum was performed similar to Snyder et al. (*23*). In brief, 1-10ng cfDNA was dephosphorylated using FastAP Thermosensitive Alkaline Phosphatase (Thermo Scientific cat. EF0651), denatured, and incubated overnight with CircLigaseII (Lucigen cat. CL9025K) and 0.093-0.125 µM biotinylated CL78 primer (*23*) at 60°C with shaking every 5 minutes. Captured cfDNA fragments were denatured and then bound to magnetic streptavidin M-280 beads (Invitrogen cat. 11205D) for 30 minutes at room temperature with nutation. Beads were washed and second-strand synthesis was performed using Bst 2.0 DNA polymerase (NEB cat. M0537) with an increasing temperature gradient 15-31°C with shaking at 1750 rpm. Beads were washed and a 3’ gap fill was performed using T4 DNA polymerase (Thermo Scientific cat. EL0011) for 30 minutes at room temperature. Beads were washed and a double-stranded adapter was ligated using T4 DNA ligase (Thermo Scientific cat. EP0062) for 2 hours at room temperature with shaking at 1750 rpm. Beads were washed and resuspended in 30 µL 10 mM TET buffer (10 mM Tris-HCl pH 8.0, 1 mM EDTA pH 8.0, 0.05% Tween-20). Beads were denatured at 95°C for 3 min and cfDNA libraries were collected after immediate magnetic separation.

Quantitative real-time PCR was performed on cfDNA libraries using iTAQ Supermix (Bio-Rad cat. 1725124) and Ct values were used to determine the number of PCR cycles needed to amplify each library. PCR was performed with KAPA HiFi DNA polymerase (Kapa Biosystems cat. KK2502) using barcoded indexing primers for Illumina. Primer dimers were removed from the libraries using AMPure beads (Beckman Coulter cat. A63881). Libraries were eluted in 0.1X TE and concentrations were determined using Qubit. The length distribution of each library was assessed by the Agilent Bioanalyzer using the D1000 or HSD1000 cassette. Libraries were sequenced for 150 cycles in paired-end mode on NovaSeq 6000 system at University of Colorado Cancer Center Genomics Shared Resource.

### CUT&RUN

We used an immuno-tethered strategy for profiling the binding of the ERα and FOXA1 transcription factor in human MCF7 breast cancer cells. MCF7 cells were estrogen withdrawn for 72 hours before being plated and then treated with either ethanol (vehicle control) or 10^−10^ M E2 (estradiol) for 1 hour prior to cell collection. The CUT&RUN method uses an antibody to a specific chromatin epitope to tether Protein A-MNase at chromosomal binding sites within permeabilized cells. The nuclease is activated by the addition of calcium and cleaves DNA around binding sites (*19*). Cleaved DNA is isolated and subjected to paired-end Illumina sequencing to map the distribution of the chromatin epitope genome-wide. We used a primary antibody to human ERα (ab3575, abcam, Cambridge, MA) and human FOXA1 (ab170933) and protein A-MNase fusion (*19*) (pA-MNase, a gift from S. Henikoff, Fred Hutchinson Cancer Research Center, Seattle WA). CUT&RUN profiling with 5×10^5^ cells and library amplification with 13 cycles of PCR was performed as described (*19*). Libraries were sequenced for 10 million paired-end reads on the Illumina NovaSeq 6000 platform at the University of Colorado Denver Cancer Center Genomics Shared Resource. Paired-end reads were mapped to the GRch38 assembly of the human genome using Bowtie2 (*67*).

### Data and code availability

All datasets were aligned to the hg38 version of the human genome. Datasets generated in this study have been deposited in GEO under accession GSE171434 and will be made public upon acceptance. All scripts and pipelines used in this study are available at https://github.com/satyanarayan-rao/tf_nucleosome_dynamics.

### CUT&RUN peaks

To call peaks, we used custom python script (deposited in github). Briefly, we first normalized coverage of <120 bp protected fragments in CUT&RUN data at 10 base pair resolution, and then smoothed the coverage with a Savitzky–Golay filter (*68*) available as a SciPy (*69*) method ‘signal.savgol_filter’ with parameters window_length = 9, polyorder = 1. We determined the cut-off for each dataset by iteratively eliminating outliers and used ‘find_peaks’ method in SciPy to call peaks that were separated by at least 250 bp.

### Aligning mouse extracted cfDNA to in silico concatenated genome

The names of chromosomes of human (hg38; GRCh38 assembly) and mouse (mm10: GRCm38 assembly) reference genomes were first prefixed by *hg38* and *mm10* respectively, and then the fasta files were concatenated together to represent an *in silico* human + mouse genome. We then aligned C/PDX cfDNA to this concatenated genome using *bowtie2* (*67*) with parameters “--local --very-sensitive-local --no-unal --no-mixed --no-discordant -I 10 -X 700”. We selected for mapped reads and then filtered out reads with secondary alignment from the bam file using the command “samtools view -F 4 <bam file> | grep -v ‘XS:’” (*70*). This filtering ensured that we did not consider any reads that aligned to both human and mouse genomes. To get human aligned reads we filtered for the *hg38* prefix in the reads’ chromosome name.

### Defining TFBSs under ChIP-seq peaks

We first selected for ChIP-seq peaks that do not overlap with ENCODE profiled blacklisted regions, and we considered all peaks except the ones on chromosome Y. We then used FIMO (*71*) with parameters “--max-stored-scores 10000000 --oc <output-directory> <motif-file> <fasta-file>“ to scan for motifs on sequences underlying ChIP-seq peaks. In case of overlapping peaks in 50 bp span, we keep the motif with higher FIMO score. Final number of motifs under ChIP-peaks used for TFs are tabulated in **Supplementary Table 2**.

### cfDNA length distribution clustering

Length distribution of mapped cfDNA fragments to a TFBS is estimated by ‘*density’* function in R with a smoothing bandwidth (bw) of 3 at 100 equally spaced points (*n* = 100) between 35 to 250 bp. Clustering of estimated cfDNA length distribution at individual sites was performed using ‘*kmeans’* function in R with parameters: *centers* = 6, *iter*.*max* = 250, and *nstart* = 20. A cluster is visually represented by the mean of fragment length distributions of sites in that cluster. Weighted length of each cluster was calculated by multiplying fragment length to its normalized frequency. Clusters 1 to 6 were assigned by ranking the clusters by their weighted length.

### Mapping cfDNA length class to TFBS and its flank

Genome-wide cfDNA read density (*bigwig*) was generated for short (< 80bp) and nucleosomal sized fragments (130-180 bp). First, a bedgraph (coverage of bases genome-wide; no normalization performed) file was generated using bedtools (*72*) genomecov utility with command line option “-bga” and then bedgraph file was converted to bigwig using kent tools “bedGraphToBigWig” (*73*). While creating the *bigwig* file we considered cfDNA fragment center ± 30bp (if fragment is >60 bp). Bigwig is mapped to TFBS±1Kb using pyBigWig module from deeptools (*74*) and then enrichment over mean (E.O.M) is calculated. E.O.M is smoothed using Savitzky–Golay filter (*68*) available as a SciPy (*69*) method ‘signal.savgol_filter’ with parameters window_length = 51, polyorder = 3.

### ChIP-seq score calculation sites in cfDNA length clusters

For a TFBS in a given cluster, *Log2* of mean fold enrichment over control was calculated for TFBS ± 300bp. pyBigWig module from deeptools (*74*) was used to map signal from bigwig file to defined genomic regions.

### MNase signal mapping to CTCF sites

MNase data from ENCODE (*18*) was mapped to CTCF motif center ± 1kb. E.O.M and smoothing was performed similar to how it was done for cfDNA length class heatmaps (see *Mapping cfDNA length class to TFBS and its flank*).

### V plots

For CTCF sites in cfDNA length clusters 1 and 2, cfDNA fragment centers were mapped to CTCF motif center ± 500 bp. Total number of cfDNA centers of a given length is plotted against the distance of the fragment centers from the CTCF motif center.

### CUT&RUN score calculation

CUT&RUN score has been calculated as the read density in regions spanning CUT&RUN peak summit ± 50 bp.

### Defining significant sites and specific sites

cfDNA length clusters that have significantly higher binding scores (ChIP scores for CTCF, PU.1 and LYL1; CUT&RUN scores for ER and FOXA1) compared to cluster 6 are considered significant i.e., overall, sites in these clusters have stronger binding strength inferred from TF binding experiments compared to cluster 6. Specific sites are identified by subtracting significant sites of one sample from significant sites from another sample. In the case of disease state detection analysis i.e., healthy vs. cancer, cancer-specific sites (CSS) and healthy-specific sites (HSS) were defined. Cancer-specific sites for ER, for example are defined by subtracting sites in healthy plasma (IH02) (*23*) significant clusters 1 and 2 from UCD65 clusters 1-4. Similarly, healthy-specific sites for ER are defined by subtracting sites from UCD65 clusters 1-4 from IH02 clusters 1 and 2. In the case of cancer state detection analysis i.e., separating tumor subtypes (UCD65 vs. MCF7, UCD4 vs. UCD65, and UCD4 vs. MCF7) using tumor TF binding sites, tumor-specific sites were defined by a similar approach. We did not observe enrichment at FOXA1 binding sites in UCD4 dataset thus tumor-specific sites were not defined for FOXA1 in UCD4.

### Dilution analysis

#### Disease detection

*In silico* patient data was generated by diluting healthy sample (IH02) (*23*) with different fractions of UCD65 cfDNA. For each dilution level, 100 *in silico* patient datasets were generated by randomly sampling reads from IH02 and UCD65 datasets at the ratio defined by the dilution level. For a given cancer/healthy-specific binding site, the TF binding score was calculated as the ratio of the short fragment coverage in (< 80bp) TFBS ± 50 to the coverage in TFBS ± 1kb. Reference TF binding score is calculated just in healthy state, and for each *in silico* patient dataset, scores are calculated in same fashion. ΔScore (used in **Figure 6C**) for cancer specific sites was calculated as the difference between patient and healthy states (gain in score), but for healthy-specific sites the sign was reversed (loss in score). T-test was performed on ΔScore values from all sites (healthy-specific + cancer-specific) to reflect how many standard deviations away the scores are from the healthy reference.

#### Cancer state detection

For each xenograft (UCD4, UCD65 and MCF7) model, 100 *in silico* patient data was generated by diluting healthy plasma (IH02) with different fractions of ctDNA. For each of three comparisons of xenograft models, the following were calculated (using UCD65 vs. MCF7 as an example): i) TF binding scores at tumor subtype specific sites using UCD65 and MCF7 *in silico* patient data respectively, ii) calculated ΔScore for UCD65-specific sites by subtracting scores of MCF7 dilution from UCD65 dilution. Similarly, ΔScore for MCF7-specific sites were calculated by subtracting scores pr UCD65 dilution from MCF7 dilution, and iii) calculated T-statistics on ΔScore using ‘ttest_1samp’ function from scipy.stats module (*69*) with expected value in null hypothesis = 0.

### TCGA ATAC-seq and expression analysis

FPKM files for each cohort were downloaded from TCGA website. FPKM for a gene was converted to TPM using the following formulae:

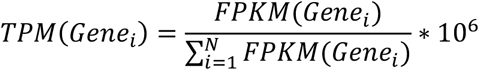

where N is the total number of genes found in the FPKM table.

ATAC insert bigwig files from Corces MR et al., (*59*) were used to map ATAC signal around TF sites (peak ± 150 bp).

### Cancer vs. Healthy and Breast Cancer vs. non-Breast cancer prediction analysis

Healthy-specific sites (HSS) and Cancer-specific sites (CSS) were ordered by their binding strength inferred from ChIP (motif center ± 300 bp; for PU.1, LYL1, and CTCF) or CUT&RUN (summit ± 100 bp; for ER and FOXA1) and grouped in a bin of size 250 to define TF features. cfDNA-inferred binding score at TF features is defined by the following formulae:

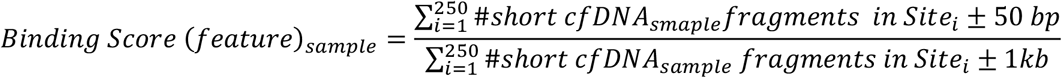

To identify what TF features are class-specific (for example, class1 – cancer, class2 – healthy), we defined a Z-score metric using the following formula:

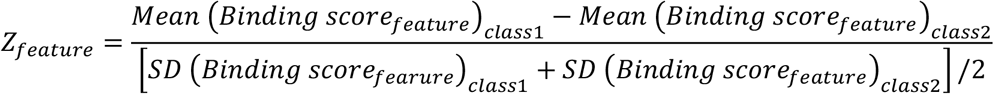

Where *SD* stands for standard deviation. Features with |*Z*_*feature*_|> 1 were selected and depending on the sign were annotated as class1-specific (+ve) or class2-specific (-ve). Enrichment of a TF in particular category (for example healthy-specific) was calculated by abundance of the TF features as log2 (Observed frequency/expected frequency).

To predict a class (breast cancer or non-breast cancer) for a cfDNA sample, leave-one-out cross validation approach was adopted where cfDNA sample of our interest was kept away during feature selection process described above. Each sample was then assigned a single score by subtracting the sum of binding scores of features with negative Z-scores (Z_feature_<-1) from the mean of features with positive Z-scores (Z_feature_>1) and then dividing by the total number of features (|*Z*_*feature*_ >1|). For the left-out sample, distances from the median of two classes were calculated and assigned the class label with closest distance.

