## Supplemental Figures and Tables for "Mapping Transcription Factor-Nucleosome Dynamics from Plasma cfDNA"

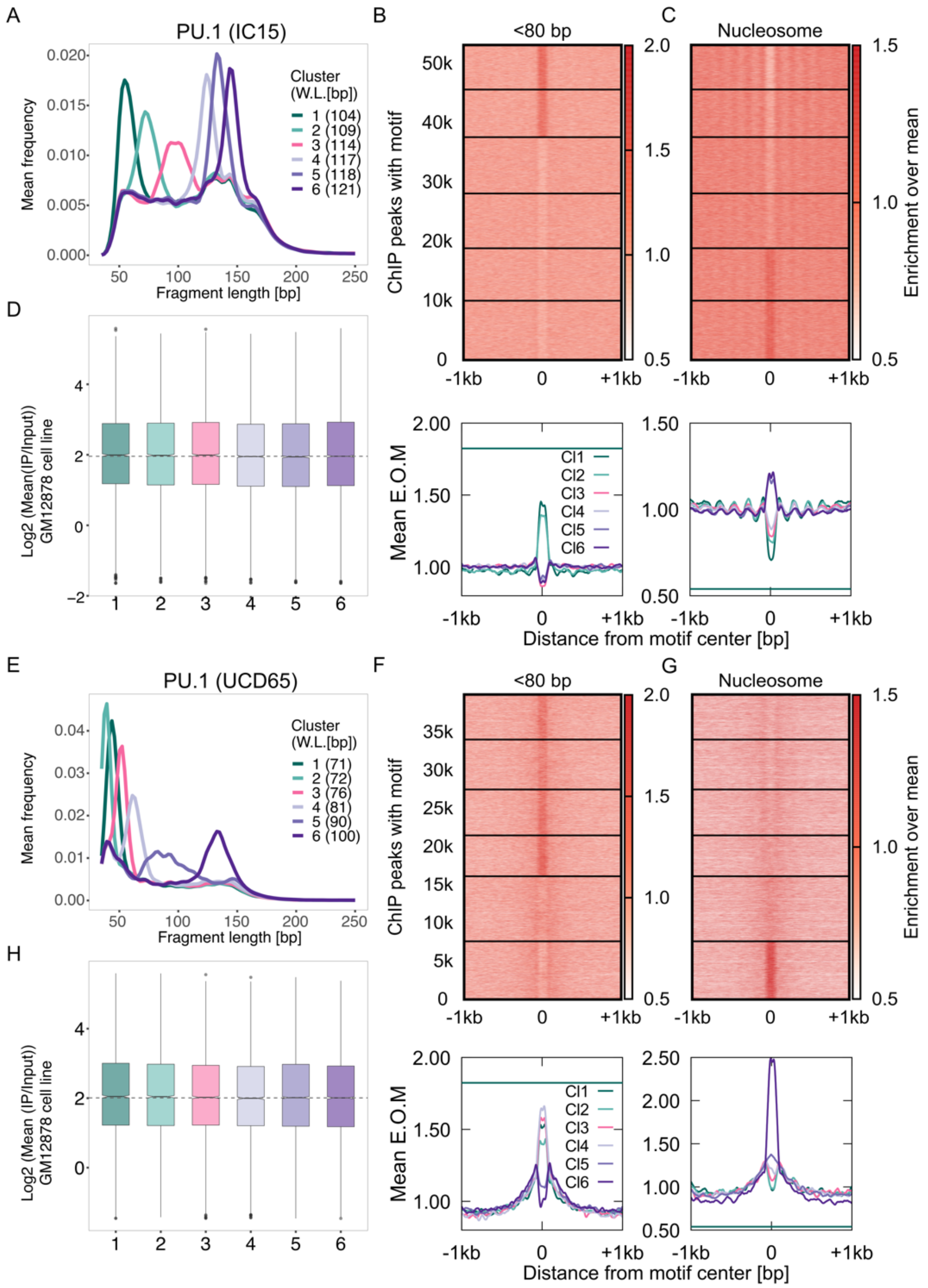


**Figure S1. Dilution of PU.1 TF footprints due to presence of cancer. A)** K-means clustering is performed on smoothed length distribution to group TFBSs with similar cfDNA fragment length distribution. Here, smoothened length distributions of clusters of PU.1 TFBS are shown. Weighted length (W.L.) for each PU.1 length cluster is shown in parentheses. **B)** Enrichment over the mean signal in TFBS ± 1Kb of cfDNA short (<80 bp) fragments is plotted as a heatmap (**top**, 53,008 PU.1 TFBS) and as metaplots for each cluster (**bottom**; the horizontal green line is drawn at the maximum mean E.O.M (enrichment over mean) observed IH02) **C)** Same as (B) for nucleosomal sized fragments. Horizontal line in the metaplot (bottom) is drawn at the minimum mean E.O.M observed in when IH02 cfDNA is mapped at PU.1 sites **D)** Mapping of ChIP scores from GM12878 cell line to length clusters. Number of sites (n) in length clusters and p-value using KS test are: Cl1: n = 7440, p (1,6) = 0.087; Cl2: n = 8032, p (2,6) = 0.58; Cl3: n = 9503, p (3,6) = 0.24; Cl4: n = 9254, p (4,6) = 0.93; Cl5: n = 8848, p (5,6) = 0.85; Cl6: n = 9931 **E-H)** Same as A-D when UCD65 ctDNA is mapped to PU.1 sites (n = 39,841). For (H) Number of sites (n) in length clusters and p-value using KS test are: Cl1: n = 5847, p (1,6) = 0.088; Cl2: n = 6537, p (2,6) = 0.093; Cl3: n = 6021, p (3,6) = 0.14; Cl4: n = 5346, p (4,6) = 0.26; Cl5: n = 8524, p (5,6) = 0.28; Cl6: n = 7566

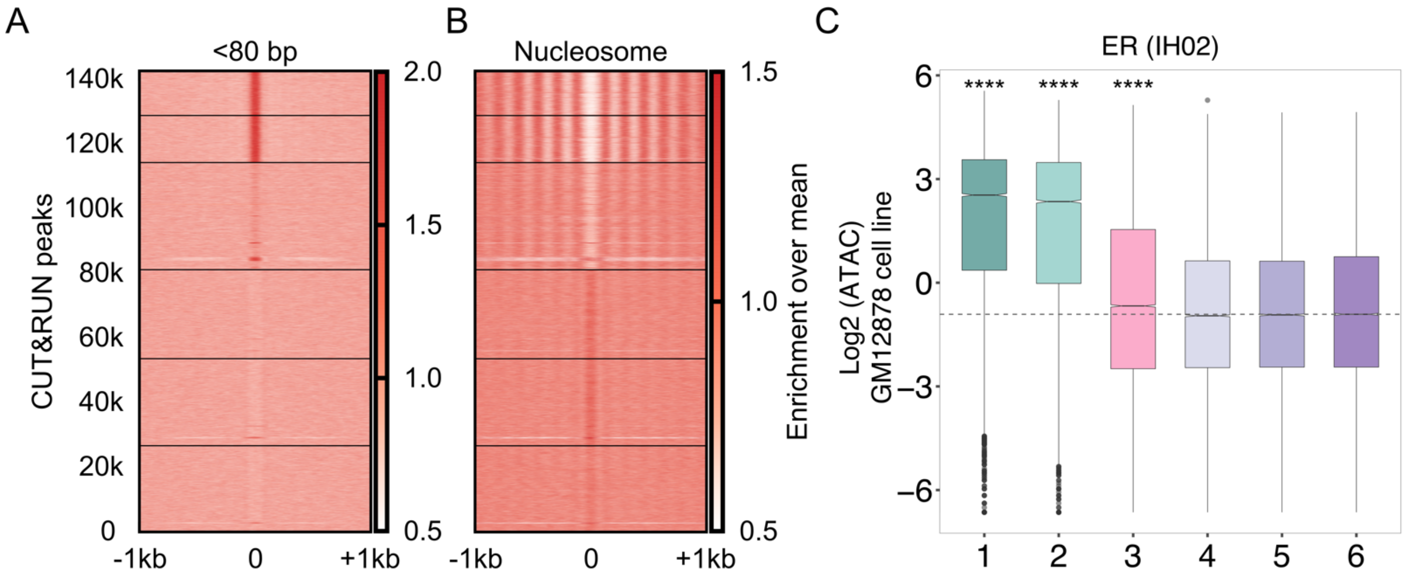


**Figure S2. ER binding inferred in hematopoietic cells**. **A)** Enrichment over the mean signal in TFBS ± 1Kb of cfDNA short (<80 bp) fragments is plotted as a heatmap (14,2202 ER CUT&RUN peaks) **B)** Same as **(A)** for nucleosomal sized (130-180 bp) fragments **C)** Boxplot of ATAC scores (Log2) from GM12878 cell line. Number of sites (n) in length clusters and p-value using KS test are: Cl1: n = 13539, p (1,6) < $2.2 \times 10^{-16}$; Cl2: n = 14586, p (2,6) < $2.2 \times 10^{-16}$; Cl3: n = 33137, p (3,6) = $1.3 \times 10^{-76}$; Cl4: n = 27534, p (4,6) = 0.23; Cl5: n = 26957, p (5,6) = 0.93; Cl6: n = 26449

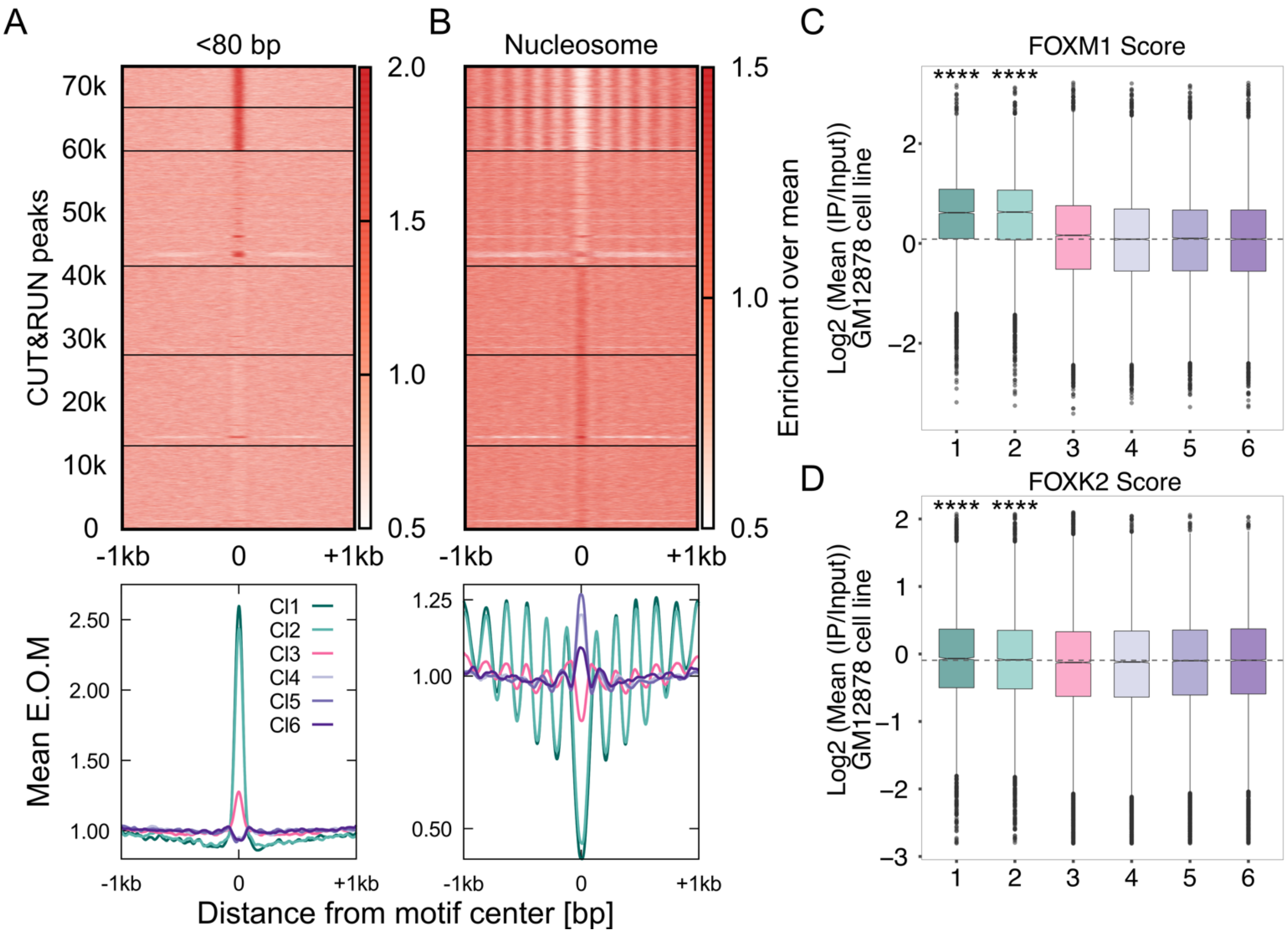


**Figure S3. FOXM1 binding in healthy plasma at FOXA1 binding sites inferred from ER+ tumor. A)** Enrichment over the mean signal in TFBS ± 1Kb of cfDNA short (<80 bp) fragments is plotted as a heatmap (**top;** 72,938 FOXA1 CUT&RUN peaks) and as a metaplot (**bottom**) **B)** Same as (A) for nucleosomal sized fragments (130-180 bp) **C)** Boxplot of FOXM1 ChIP scores from GM12878 cell line mapped to cfDNA length clusters. Number of sites (n) in length clusters and p-value using KS test are: Cl1: n = 6351, p (1,6) = 2.4 x$10^{-290}$; Cl2: n = 6871, p (2,6) = 6 x$10^{-259}$; Cl3: n = 18216, p (3,6) = 0.01; Cl4: n = 14075, p (4,6) = 0.23; Cl5: n = 14375, p (5,6) = 0.92; Cl6: n = 13050. **D)** Boxplot of FOXK2 ChIP scores from GM12878 cell line mapped to cfDNA length clusters. n and p are: Cl1: n = 6351, p (1,6) = 1.2 x$10^{-20}$; Cl2: n = 6871, p (2,6) = 2.2 x$10^{-11}$; Cl3: n = 18216, p (3,6) = 0.46; Cl4: n = 14075, p (4,6) = 0.52; Cl5: n = 14375, p (5,6) = 1; Cl6: n = 13050

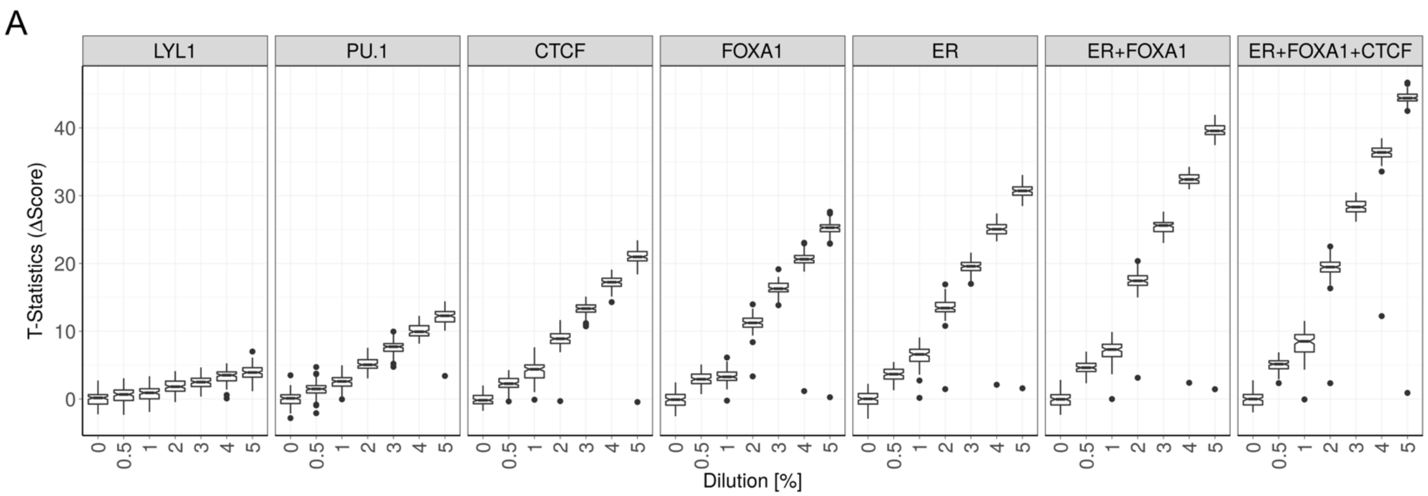


**Figure S4. Boxplots of T-statistics performed on ΔScore at different TF sites (individual or in combination) for 100 bootstraps at a given dilution.**

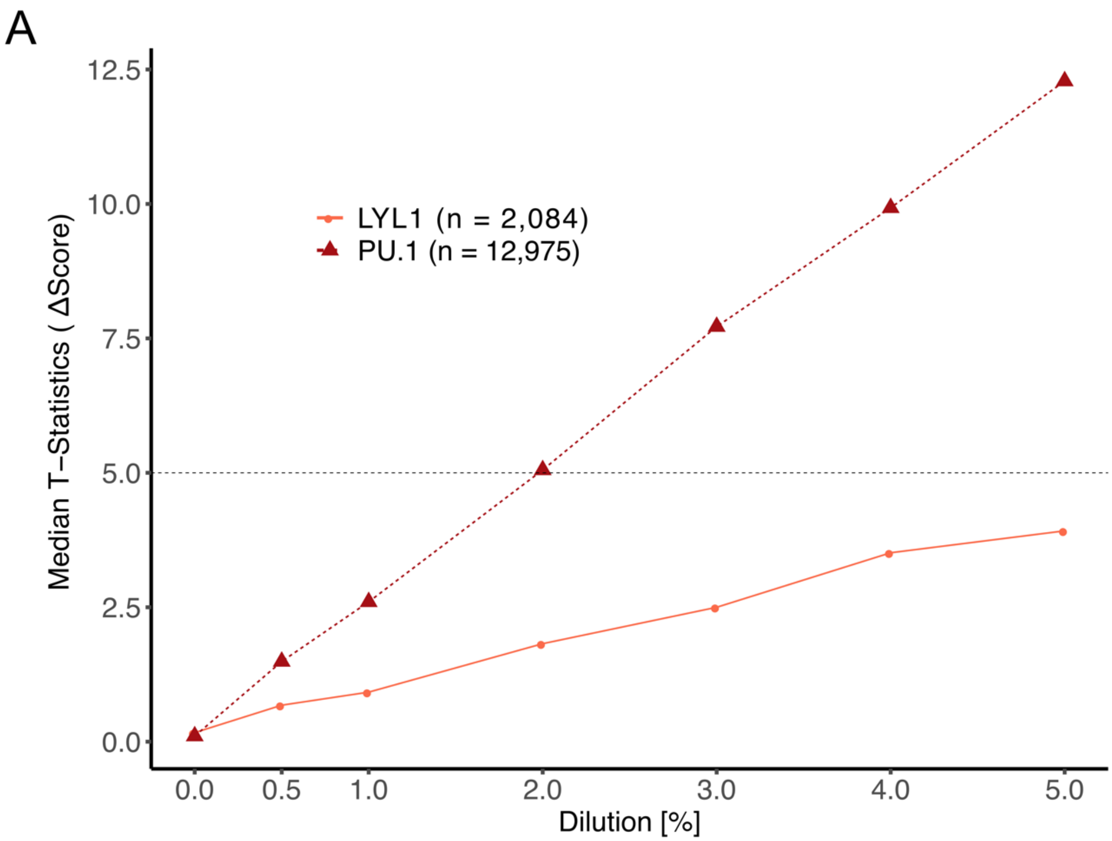


**Figure S5. Median t-statistics line plot for PU.1 and LYL1.**

**Table S1. Plasma samples used in this study.**

| Sample name | Source name | Sex | Disease status |
|---:|---:|---:|---:|
| MCF7 | Cell line | F | ER+ breast cancer |
| UCD4 | Breast tumor xenograft | F | Breast cancer with ER mutation |
| UCD65 | Breast tumor xenograft | F | Breast cancer with ER amplification |
| F02 | Cell-free DNA | F | Healthy |
| F05 | Cell-free DNA | F | Healthy |
| SporeD3 | Cell-free DNA | M | Healthy |
| BC02 | Cell-free DNA | F | ER+ breast cancer |
| BC03 | Cell-free DNA | F | ER+ breast cancer |
| SporeA2 | Cell-free DNA | M | Lung cancer (non-small) |
| SporeB2 | Cell-free DNA | F | Lung cancer (small cell) |
| SporeF2 | Cell-free DNA | M | Lung cancer (squamous) |
| SporeG2 | Cell-free DNA | M | Lung cancer (Adenocarcinoma) |

**Table S2. Transcription Factor ChIP-seq Peak Counts.**

| Category | CTCF | PU.1 | LYL1 |
|---|---|---|---|
| Total ChIP peaks | 231,309 | 67,558 | 33,709 |
| Chr1-X | 231,075 | 67,502 | 33,681 |
| After blacklisted region filtering | 230,965 | 67,496 | 33,608 |
| Motif discovery | 215,818 | 71,205 | 14,899 |
| Overlapping motifs in +- 50 bp | 97,229 | 17,216 | 6,853 |
| Non-overlapping motif | **118,589** | **53,989** | **8,046** |
